# ULTRA-Effective Labeling of Repetitive Genomic Sequence

**DOI:** 10.1101/2024.06.03.597269

**Authors:** Daniel R. Olson, Travis J. Wheeler

## Abstract

In the age of long read sequencing, genomics researchers now have access to accurate repetitive DNA sequence (including satellites) that, due to the limitations of short read sequencing, could previously be observed only as unmappable fragments. Tools that annotate repetitive sequence are now more important than ever, so that we can better understand newly uncovered repetitive sequences, and also so that we can mitigate errors in bioinformatic software caused by those repetitive sequences. To that end, we introduce the 1.0 release of our tool for identifying and annotating locally-repetitive sequence, *ULTRA* (**U**LTRA **L**ocates **T**andemly **R**epetitive **A**reas). *ULTRA* is fast enough to use as part of an efficient annotation pipeline, produces state-of-the-art reliable coverage of repetitive regions containing many mutations, and provides interpretable statistics and labels for repetitive regions. It released under an open license, and available for download at https://github.com/TravisWheelerLab/ULTRA.

## Introduction

### Background

#### Tandem Repeats

Biological sequences are complex and full of subtle patterns; however, one of the most common patterns is not subtle at all: *tandem repeats*. Tandem repeats are a class of biological sequences that, for various reasons [1–4], consist of multiple adjacent copies of some core repetitive unit. “AACAACAACAACAACAACAACAAC” is an example of a short tandem repeat composed of an “AAC” unit repeated 8 times. Tandem repeats are easy to identify when their repetitive pattern is well conserved, but as a repetitive region ages and accrues mutations, the clarity of its repetitive pattern decays. For instance, “AACAACAATATCAATAACAACAACAGCAAC” (found by *ULTRA* at location 61,461 in T2T-CHM13v2.0 chromosome 1) was likely a once pristine “AAC” repeat, but after only a few substitutions the repetitive pattern has become less obvious. Insertions and deletions further obfuscate repetitive patterns, and old tandem repeats that have experienced many mutations are frequently difficult or impossible to annotate with high confidence.

An assortment of repeat expansion mechanisms is responsible for the various classes of tandem repeats. *Short tandem repeats* [5] are primarily caused by replication slippage [1, 6] and have small repetitive units from 1 to 6 bp, which can repeat to span up to hundreds of bp [7, 8]. *Minisatellites* [9] are a larger sort of tandem repeat, having units between 6-60 bp and occupying regions as large as 20 kb [10]. *Satellite repeats* [11] are found in heterochromatin and can be millions of base pairs in length, having repetitive periods that range from a few bases to a few thousand bases [12]. *Higher order repeats* [13] are a subclass of satellite repeats that have repetitive units that are themselves composed complex and shifting patterns of satellite repeats [12].

Tandem repeats of all sorts have long been studied for reasons of scientific intrigue and also due to their effects on human health. As a brief (and incomplete) survey: they contribute to protein and RNA function [14–16], they are involved in gene regulation [17, 18], influence the evolution and maintenance of centromeres and telomeres [19, 20], and they play a role ingenetic diseases [14, 21–23].

#### Bioinformatic Challenges Caused by Tandem Repeats

Historically, locally-repetitive regions of DNA caused problems with assembly [24, 25] because they are often longer than the sequence reads produced by early generations of sequencers [26]. This challenge has been largely resolved thanks to advances in long read sequencing [27–29], which produces reads that are long and accurate enough to enable assembly of centromeric satellite DNA, as exemplified by the first “complete” human reference genome, T2T-CHM13 [30, 31].

Despite these gains, tandem repeats still pose substantial problems for bioinformatic analysis. Consider homology search with tools like *BLAST* [32] and *HMMER* [33]; these tools provide a means to estimate the probability that two sequences are evolutionarily related to one another (i.e homologous). Briefly, *BLAST* and *HMMER* work by measuring the similarity between two sequences and then estimating how often that level of similarity is expected to occur by random chance [34]. Tandem repeats occur far more frequently than expected by the simplistic models of random chance used by homology search tools, and as a result, tandem repeats are a frequent source of false positive search results. For example, consider the short tandem repeat “ATATATATATATATATATATATATATATATAT”. With naive notions of random chance we would expect that exact sequence to occur once every 1.8 × 10^17^ nucleotides (assuming 60% AT richness), corresponding to a 1 in 10^7^ chance of the sequence being found in the human genome. However, in human (T2T-CHM13v2 [30]) chromosome 1, there are 624 non overlapping occurrences of that exact tandem repeat (as found with “`grep -oi`”), and 3998 non overlapping occurrences of the tandem tandem repeat with 5 or fewer substitutions (found by running “`ugrep -oi -Z` ∼`5`” on T2T-CHM13v2 Chr 1 with newlines removed). The majority of these “AT” repeats are not similar to one another because of homology, but because of independent replication slippage events that happen to produce similar short tandem repeats. Regardless of true homology, when a query sequence that contains the “AT” repeat from above is searched against human chromosome 1, that search will yield false positive hits for each non homologous “AT” repeat. The problem is not limited to sequences that contain perfect or near perfect tandem repeats – even highly decayed tandem repeats can and do induce high scoring false positive search results [35].

#### Reducing Repeat-Caused Error

The most common way to alleviate bioinformatic error caused by tandem repeats is through masking [36, 37]. A strategy called *hard-masking* uses a tandem repeat annotation tool to find and then mask (hide) tandem repeats from all downstream analysis by replacing the repetitive letters with the ambiguous letter N (for DNA) or X (for protein).

An alternative approach, called *soft-masking*, aims to improve sensitivity relative to hard-masking. Under soft-masking, repetitive sequence isn’t hidden completely but is instead marked as repetitive, typically by making repetitive sequence lower-case in the representative sequence file. Some homology search tools ignore soft-masked regions during the phase of identifying alignment seeds, then allow repetitive sequence to be used while creating a final sequence alignment [36].

The problem with both hard-masking and soft-masking is that one must choose some threshold of repetitiveness (i.e repetitive annotation score) to decide what should be masked and what should not be masked. Low thresholds mask too much and result in reduced annotation sensitivity [38]. High thresholds mask too little, allowing low-scoring decayed repeats to cause false positive hits [35]. One potential strategy for reducing errors caused by tandem repeats is to not mask at all, but to instead better incorporate models of repetitiveness directly into the annotation process [39]. Such an approach could proceed by either directly including repetitiveness in the process of identifying and scoring of alignment, or by competing repeat annotations against alignment based annotations [40]. Both of these approaches demand probabilistic models that produce interpretable scoring statistics.

### Repeat Annotation Tools

There is a plethora of tools used to find tandem repeats [41–49] each tool with a unique set of strengths and weaknesses. Throughout this paper we will compare our tool, *ULTRA*, against two tools in particular: *TRF* [48] and *tantan* [49].

The venerable *TRF* is the most widely used de novo repeat finding tool. *TRF* first searches for well conserved tandem repeat candidates, and then proceeds with a self-alignment algorithm that further extends the repeat annotation while also producing informative annotation labels describing the repetitive pattern and all mutations that occur throughout each observed tandem repeat.

*tantan* is based on a hidden Markov model (HMM) for repetitive sequence, and employs the Forward-Backward algorithm [50] to assign a numeric value to each letter in the target sequence representing the probability of the letter being a part of a tandem repeat. *tantan* is blazingly fast and can be tuned to provide greater sensitivity and specificity compared to *TRF*. Due to its design focus on the repetitive nature of individual letters, it does not aim to assign scores or labels to repetitive regions, and thus produces less descriptive repeat annotations than does *TRF*.

In an earlier conference paper, we described a prototype implementation of our HMM-based repeat annotation tool, *ULTRA* (**U**LTRA **L**ocates **T**andemly **R**epetitive **A**reas) [51]. Our goal in developing *ULTRA* is that it will improve sensitivity and specificity relative to other tools (particularly in the context of high mutational load), provide statistically meaningful annotation scores, create consistent and interpretable characterization of repetitive patterns, be easily re-parameterized for specific genomes, and provide a stable and easy user experience. Here, we introduce the user-ready 1.0 release of *ULTRA* and demonstrate its repeat labeling and scoring output. We also compare its performance to *TRF* and *tantan*, and show that it produces high genome coverage with low false labeling rate.

## Methods

### ULTRA’s Hidden Markov Model

#### Repetitive and Nonrepetitive States

*ULTRA* models tandem repeats using a hidden Markov model (HMM – for an introduction to HMMs, see [52, 53]). *ULTRA*’s HMM (Figure 1) uses a single state to represent non-repetitive sequence, and a collection of repetitive states that each model different repetitive periodicities. The non-repetitive state is blind to context and its emission distribution approximately represents how frequently letters are expected to occur in non-repetitive background sequence. In contrast, each repetitive state is *context-sensitive* [54], meaning that the repetitive emission distribution depends on prior observations. A period *p* repetitive state at position *t* has a greater chance of repeating the letter that was observed at position *t* − *p* than emitting a mismatching (non-repetitive) letter. Importantly, the repetitive emission distribution only depends on the individual letter at position *t* − *p* and is otherwise unaware of context.

**Figure 1.**
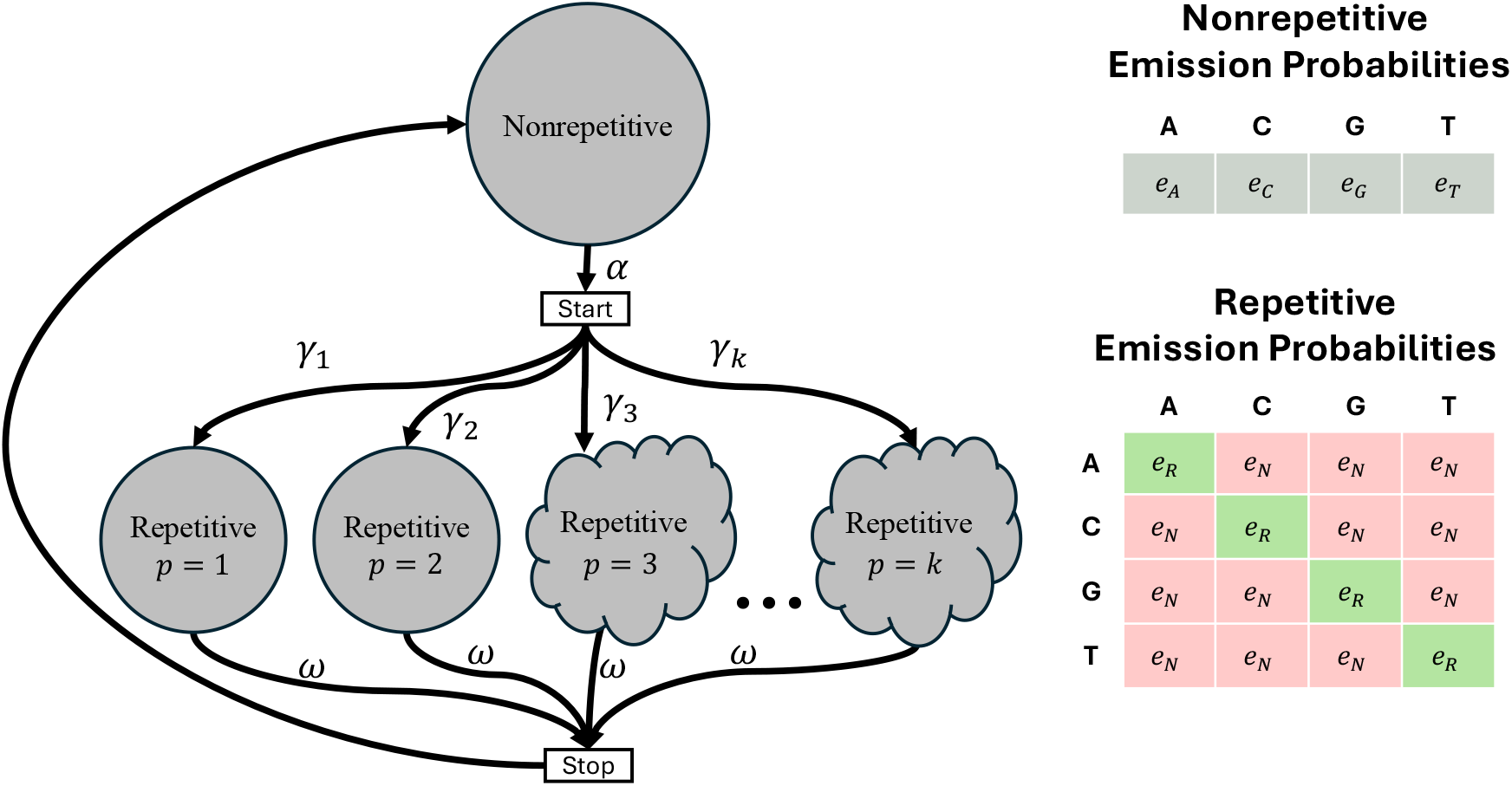
*ULTRA*’s HMM. The cloud shaped nodes represent a collection of states modeling both tandem repeats and also insertion/deletion events (see Section Insertion and Deletion States). Self-transition edges have not been drawn but do exist for the non-repetitive and repetitive states. Similar to *tantan, ULTRA* models large period repeats as being less common than small period repeats through a decay parameter, *λ*. For a model allowing maximum period k, the probability of transitioning from the *start* state to a period *p* repetitive state is 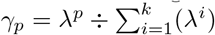. All labeled parameters can be adjusted (see userguide at https://github.com/TravisWheelerLab/ULTRA).

#### Insertion and Deletion States

Substitution mutations are modeled by the emission distribution of repetitive states. An individual substitution will cause a slight decrease in annotation score, but because it does not affect the overall repetitive pattern, it will not dramatically impact the overall repeat annotation. On the other hand, insertions and deletions (indels) cause a temporary offset in repetitive pattern and can result in a large reduction in repeat annotation score that may be significant enough to grossly change the overall repeat annotation. Consider the following example:

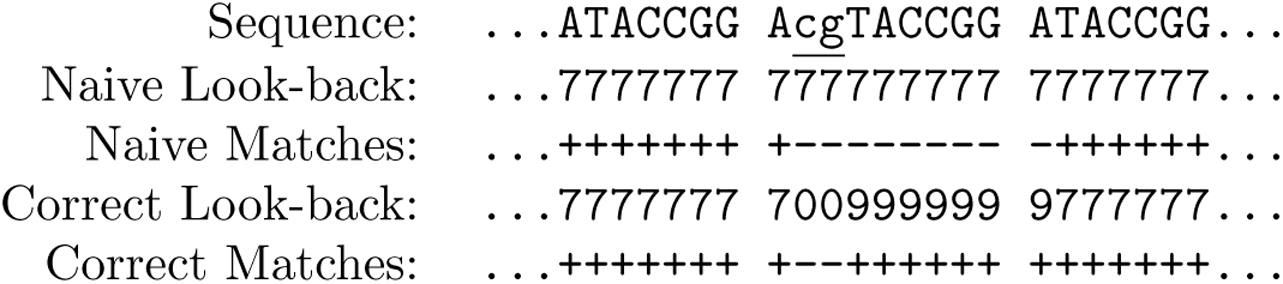

The inserted letters “cg” are effectively nonrepetitive (i.e period 0) because they are independent of the repetitive pattern. The insertion also causes a temporary offset in repetitive pattern, and if the indel is not accounted for, then the period *p* repetitive state will accumulate as many as *p* undeserved mismatches.

*ULTRA*’s HMM models indel events with period-specific indel states that account not only for inserted and deleted letters, but also for the temporary offset in repetitive pattern caused by indels (see Figure 2). *ULTRA* uses three types of states to model indels. I-states represent insertion events and are modeled as *emitting* a letter with disregard for the current repeat pattern. D-states are silent states that are modeled as *omitting* a letter in the repeat pattern. *ULTRA* uses J-states to model the offset that occurs for the *p* letters that follow an indel. More specifically, each I-state and D-state connects to a unique chain of *p* J-states that lead from the I-state or D-state back to the original period *p* repetitive state. During repeat annotation, an entire J-state chain can be updated efficiently without computing or storing values for individual J-states; this optimization is akin to updating a moving average based on which values have changed instead of simply recomputing the entire average. For a more complete description of this optimization see our earlier conference extended abstract [51]. (Note that *ULTRA*’s HMM is similar to *tantan*’s HMM, supplemented by these additional states to explicitly model the effects of insertions and deletions).

**Figure 2.**
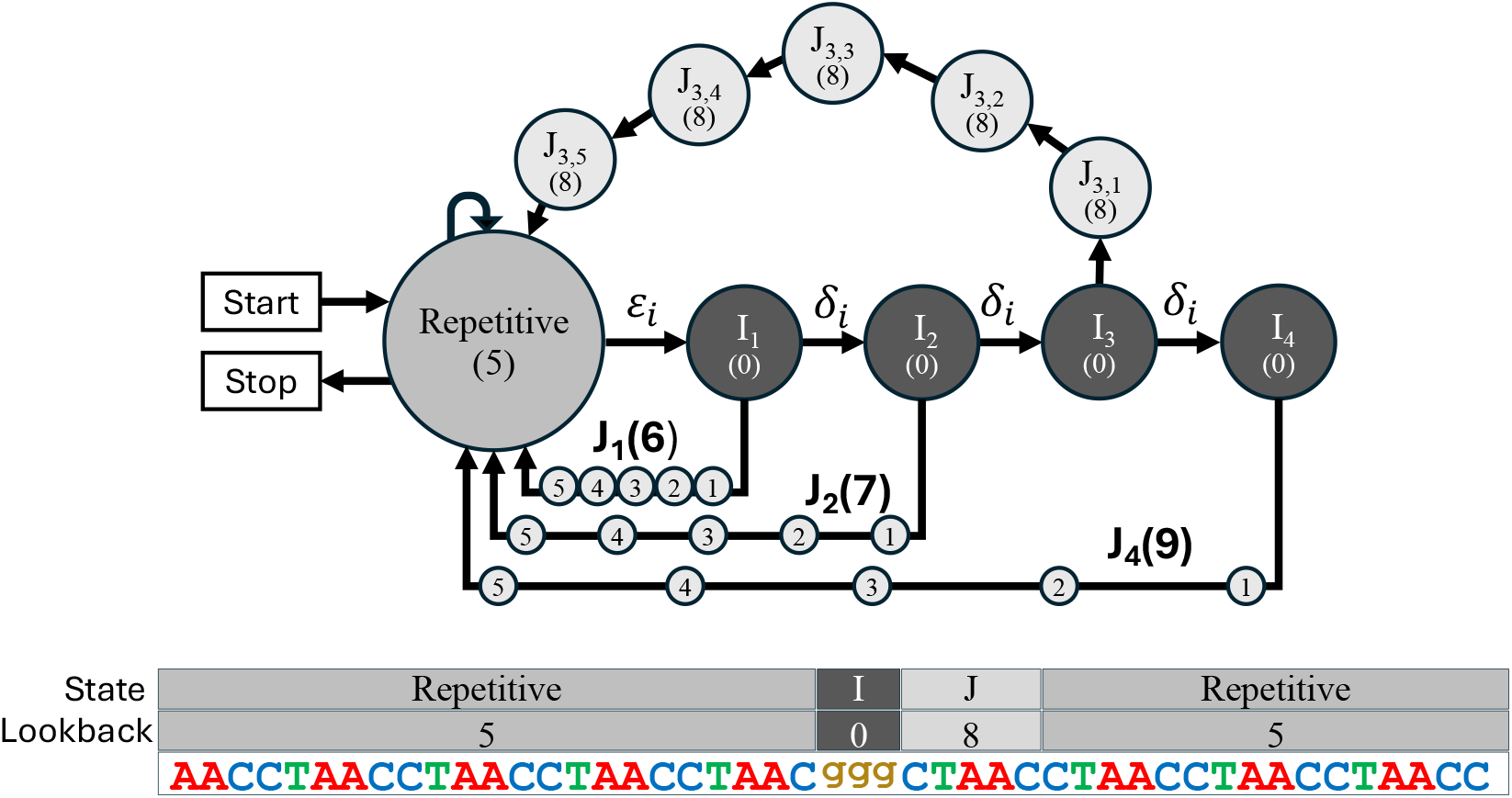
Top: The collection of states used to model insertions that occur within a *p* = 5 tandem repeat (a similar collection of states is used to model deletions). Each state’s look-back is shown within parentheses. Bottom: A *p* = 5 tandem repeat that contains a length 3 insertion. The letters from the insertion are explained by a path through 3 I-states with look-back = (0), followed by a chain of J-states with look-back = 3 + 5 = (8).

Modeling all possible patterns of indel events while keeping track of the temporary offset in repetitive pattern caused by those events would result in an intractably large HMM. Instead of modeling all possible patterns, *ULTRA* only models up to *i* consecutive insertions and up to *d* consecutive deletions where, by default, *i* = *d* = 10. Anecdotally, we find that more complex indel patterns can frequently be well approximated by *ULTRA*’s basic model of consecutive indels, and empirically we find that modeling consecutive indels provides improved annotation performance relative to modeling no indels at all.

### Viterbi Annotation

#### Basic Viterbi

To annotate repeats in a length *n* string, *S, ULTRA* finds regions of *S* that are well represented by repetitive states and then marks those regions as being repetitive. The problem of finding the sequence of HMM states with the greatest probability of producing an observed string is known as the *decoding problem*, and *ULTRA* solves that problem using the*Viterbi algorithm* [55]. Let ∑ be the model alphabet (e.g in DNA ∑ = {A,C,G,T}), *M* be the collection of *m ULTRA* HMM states (including non-repetitive, repetitive, I, D, and J states), *T* ∈ *R*^*m*× *m*^ be the transition probabilities, and 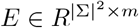 be the emission probabilities; we will use the shorthand *E*_*t,v*_ to refer to the probability of state *v* emitting the letter *S*_*t*_ ∈ ∑ at index *t*. Viterbi works by creating a matrix, *P* ∈ *R*^*n*× *m*^, that is filled using the following recurrence:

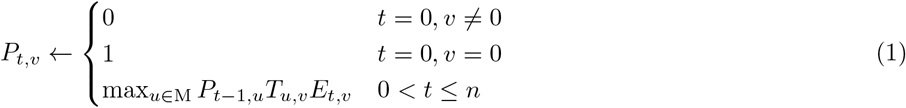

Let *v*^*\**^ be state with the greatest value in *P*_*n*_, i.e *v*^*\**^ = argmax_*u*∈*M*_ *P*_*n,u*_. The probability inside *P*_*n,v\**_ will be the probability of a most likely path through *M*, i.e the *Viterbi path*. The Viterbi path can then be recovered by re-tracing the transitions which led to *v*^*\**^ at index *n* (see [53] for a more complete description of Viterbi).

#### ULTRA Viterbi

*ULTRA*’s implementation of Viterbi replaces emission probabilities with the ratio of model emission probability relative to the background frequency of letters, *E*_0_. We refer to *ULTRA*’s Viterbi matrix as *P* ^*′*^.

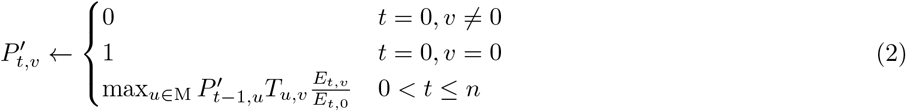

The effects of using relative emission likelihood are greatest in highly biased sequence composition, and enable *ULTRA* to better differentiate repeating letters that occur by random chance (because the letters are very common) from repeating letters that are part of a tandem repeat. Importantly, *ULTRA*’s usage of relative emission likelihood inside of Viterbi produces very different results from using basic Viterbi followed by a composition-bias adjustment; in testing we observed that using relative likelihood of emissions greatly improved *ULTRA*’s overall sensitivity and specificity.

One downside to using relative emission likelihood is that it is more difficult to predict how adjustments to the background emission rate parameters will affect repeat annotation. In future updates to *ULTRA* we wish to improve the interpretability of *ULTRA*’s background probability rate, but until then we suggest that users take advantage of the `--tune` subprogram (described in Section Tuning), which automatically optimizes *ULTRA*’s emission probabilities for a given input sequence.

#### Using the Viterbi Path to Annotate Repeats

After the Viterbi path, *V* = {*v*_0_, *v*_1_, …, *v*_*n*_}, has been calculated it is then used to annotate tandem repeats. A substring with range (*t*_start_, *t*_end_) will be annotated as repetitive when all of the positions in the range are identified by the Viterbi path as having been emitted by a repetitive state (or its associated indel states). In other words, a region is labeled as repetitive when *∀t* ∈ {*t*_start_, …, *t*_end_}, *v*_*t*_ ≠. After a *p* period repeat has been found in the range (*t*_start_, *t*_end_), it is then scored using the following equation:

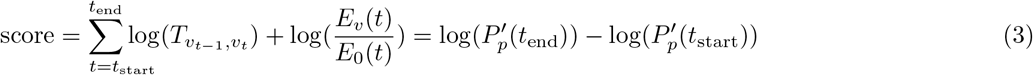

Because repetitive states in *ULTRA* look backwards but not forwards, the first *p* letters of a repeat will not be part of the Viterbi path. *ULTRA* remedies this problem by reporting the starting location for repeats as *t*_start_ − *p* instead of *t*_start_.

### Repeat Splitting

“ACACACACACACGCGCGCGCGCGCGC” is a tandem repeat that changes from an “AC’ pattern to a “GC” pattern; because repetitive states in *ULTRA*’s repeat model only compare the letter at position *t* to the letter at position *t* − *p, ULTRA* only observes a single mismatch when the pattern changes and is therefore unaware of the change in pattern. To annotate pattern changes such as this, *ULTRA* performs a post processing *repeat splitting* step. We refer to locations where a change in pattern occurs as *repeat splits*, and we refer to the segments of repetitive sequence isolated by repeat splits as *subrepeats*.

*ULTRA*’s repeat splitting process (see Figure 3) begins by assigning *profile-indices* to letters in the sequence. The profile-indices are used to keep track of how letters in the repetitive sequence relate to the repetitive pattern. For example, the sequence “CATCATCAgTCATCAT” is assigned the indices “**1** 2 3 **1** 2 3 **1** 2 * 3 **1** 2 3 **1** 2 3”. *ULTRA* then slides two adjacent windows along the tandem repeat and uses the sequence and profile indices to build window-specific repeat-profiles, *L* and *R*. The profiles are stored as matrices; a period *p* repeat with an alphabet ∑ will have a corresponding profile matrix of size *p* × |∑|. Each *L*_*i,c*_ and *R*_*i,c*_ cell holds the frequency with which letter *c* is observed at profile-index *i* in the window. The repetitive content of *L* is then compared against the repetitive content of *R* by calculating the Jensen-Shannon divergence [56, 57]:

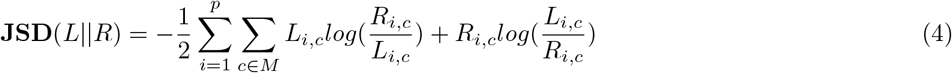

**Figure 3.**
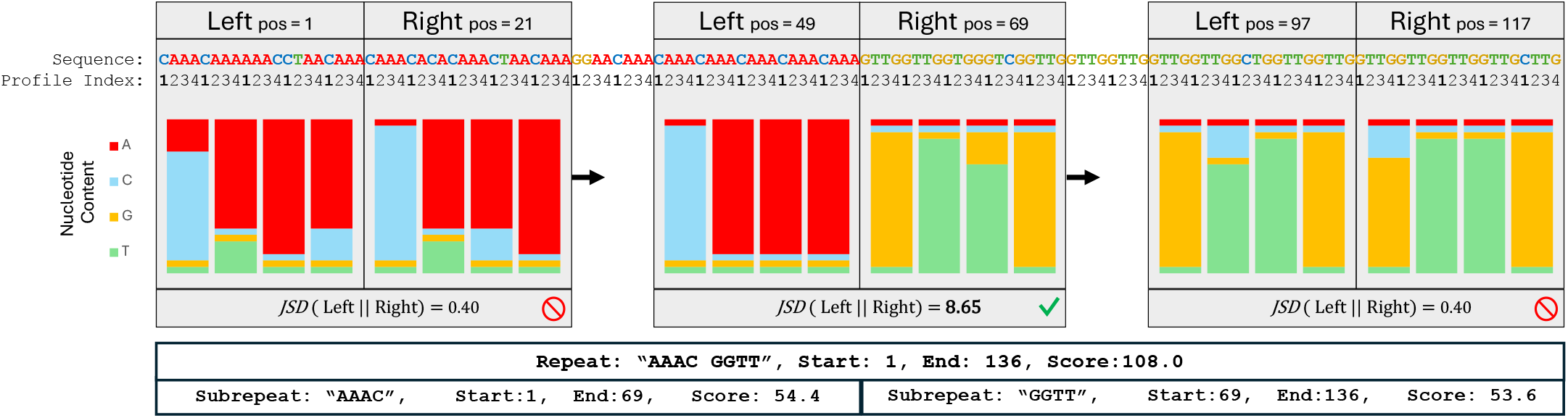
A period 4 repeat with two subrepeats (“AAAC” and “GGTT”), each containing multiple substitutions. To find the change in pattern, *ULTRA* slides two adjacent windows along the sequence and creates profiles representing the repetitive content within the windows. The repetitive profiles of two adjacent windows are compared against each other using JSD. This figure shows the local window profiles (with profile frequencies displayed as bar charts) and the corresponding JSD for three different positions. The first pair of window profiles contains similar repetitive content resulting in a small JSD; the last profile pair also yields a small JSD. The middle windows contain different repetitive content, resulting in a large JSD that passes *ULTRA*’s splitting threshold. Both the repetitive region as a whole and also the repetitive region’s subrepeats are included in *ULTRA*’s final annotation.

*ULTRA* slides the two repeat profile windows across the entire repetitive sequence and iteratively calculates the JSD at each position in the repetitive sequence. A small JSD implies that both windows contain similar repetitive content and can be well represented as a single repetitive region. Alternatively, a large JSD means that the left and right windows contain dissimilar repetitive content. ULTRA identifies local peaks in JSD, and when the value of a peak exceeds some threshold (controlled by `--split val`), the corresponding position is marked as a repeat split. After splits have been marked, the repetitive patterns are calculated for each subrepeat.

In cases where an undetected indel has occurred, the profile indices will be incorrectly assigned resulting in a large JSD even when the repetitive pattern hasn’t actually changed. *ULTRA* accounts for this possibility by comparing neighboring subrepeats to ensure the subrepeats have different repetitive patterns. When two neighboring subrepeats have the same (possibly rotated) repetitive pattern (such as ‘GTTG’ and ‘GGTT’), the repeat split is removed and the two subrepeats are merged back into a single subrepeat. The final annotations output by *ULTRA* describe the overall repetitive region as well as the subrepeats that make up the region. Repeats and subrepeats are named with an alphabetically sorted version of their repetitive pattern (for example, the pattern ‘GGTT’ from Figure 3 is alphabetically smaller than rotations ‘GTTG’, ‘TTGG’, and ‘TGGT’).

### Calculating P-values

It is possible to base repeat annotations directly on scores, by establishing some score threshold *τ* and identifying a candidate region as repetitive if its *ULTRA* score exceeds *τ*. But sequence annotation is more robust when scores can be interpreted in a likelihood framework; this is common for alignment-based annotation, where each alignment with score *s* is assigned a P-value (and multiple-test-adjusted E-value) that corresponds to the probability of observing a score ≥ *s* in an alignment of two unrelated sequences. *ULTRA* similarly computes a P-value for each annotated region by learning the distribution of scores observed in random (non-repetitive) sequence.

*ULTRA* was used (with default settings) to annotate repetitive regions in 1 Gb of randomly generated 50% AT-rich sequence. We then fit an exponential distribution parameters (location = *μ*, and scale = *σ*) to the distribution of annotation scores observed in the 1 Gb random sequence. Letting *ω* be the fraction of letters annotated as repetitive, and *s* be the *ULTRA* annotation score of a labeled repetitive region, the P-value of that region can be calculated as `P-value`(`score`) = *ω* exp ((*μ* − *s*)*/σ*).

The `--pval` flag will instruct *ULTRA* to convert annotation scores to P-values. We note that changes in *ULTRA’s* model parameters or changes in sequence composition can greatly effect the location, scale, and repeat frequency parameters that should be used to produce P-values. In a future release, we plan to automatically extract location, scale, and repeat frequency during tuning (see Section Tuning). Until then, users may modify P-value parameters with the `--ploc`, `--pscale`, and `--pfreq` options.

### Windowed Viterbi

Applying Viterbi to a length *n* sequence using an HMM with *m* states requires storing values in an *n* × *m* matrix. The size of that matrix can be a limiting factor when analyzing large sequences with large models. To reduce the amount of memory required, *ULTRA* splits an input sequence into overlapping windows and then applies Viterbi to each window independently. Dividing the target sequence into windows has the added benefit of enabling parallel computing of repeats across distinct windows. Using overlapping windows introduces the possibility of tandem repeats that occur in the overlapping region of windows being annotated multiple times. To avoid redundant repeat annotation, *ULTRA* will search for and then merge overlapping repeats when they are equal in repetitive period.

*ULTRA* automatically adjusts the size of sequence windows and the amount of overlap between windows based on the number of states *ULTRA*’s HMM is using; when using a large collection of states the window size is decreased (in order to reduce memory usage) and when using a small collection of states the window size is increased (to improve runtime). Window size and overlap size can also be manually adjusted with the `--winsize` and `--overlap` options. Users can see how much memory *ULTRA* is expected to consume by using the `--mem` flag.

### Tuning

There is tremendous diversity in tandem repeat patterns between different organisms. To achieve the highest annotation quality, *ULTRA*’s parameters need to be tuned according to the genome being annotated. In *ULTRA*, parameter tuning is performed automatically when used with the `--tune` flag. Tuning works by running *ULTRA* many times on the input sequence, each time using a different set of candidate parameters. *ULTRA* then selects the parameter set that provides the greatest annotation coverage while remaining under an estimated false discovery rate threshold (10% by default).

To estimate the false discovery rate of a particular parameter set, *ULTRA*’s tuning routine measures *ULTRA*’s annotation coverage on a locally shuffled version of the input sequence. *ULTRA*’s local shuffling is done inside the sequence windows used by windowed Viterbi (see Section Windowed Viterbi). Local shuffling retains local variability in sequence composition, as is observed in isochores [58–60], but removes all biologically caused tandem repeats. By random chance, the shuffled sequence will still contain some repetitive content, but tandem repeats that occur in shuffled sequence are expected to be rare and low-scoring relative to the original (biological) input sequence. After *ULTRA* has used candidate parameters to annotate both the input sequence and the shuffled input sequence, it then estimates the false discovery rate as repetitive coverage in the shuffled sequence divided by the repetitive coverage in the original input sequence.

*ULTRA* allows users to change what parameter configurations will be tested during tuning. `--tune` explores 18 different parameter configurations, `--tune medium` explores 40, and `--tune large` explores 252. Users can also provide their own candidate parameter configurations using the `--tune file` option. By default, *ULTRA* reduces the runtime of tuning by disabling indel states. After indel-free identification of good parameters, *ULTRA* is re-run with indels enabled. This approach can sometimes lead to selection of parameters that produce higher-than-expected false discovery rate once the indel model is enabled in the final run; to bypass this risk, indel states can be enabled during tuning with the `--tune indel` flag.

## Results

### Coverage

We quantify the sensitivity and specificity of *ULTRA, TRF*, and *tantan* by measuring each tool’s coverage and estimated false coverage for the genomes listed in Table 1. In order to estimate false coverage, we ran each tool on a window-shuffled version of each genome. Specifically, we used the *esl-shuffle* tool from *HMMER*’s [33] easel library, applying the “`-w` 20000” flag to shuffle 20kb chunks so that regional GC compositional variability would be retained while also removing (through shuffling) biologically-caused tandem repeats. We use annotation coverage of the shuffled genomes to estimate the false coverage of the unshuffled genomes.

**Table 1.** Genomes used for coverage experiments.

| Organism | Strain/Version | GC % |
| --- | --- | --- |
| *Mycobacterium tuberculosis* | H37Rv (AL123456.2) [61] | 65.6% |
| *Neurospora crassa* | NC12 [62] | 48.2% |
| *Zea Maize* | B73-REFERENCE-NAM-5.0 [63] | 46.7% |
| *Homo sapien* | T2T-CHM13v2.0 [30] | 40.8% |
| *Plasmodium falciparum* | ASM276v2 [64] | 19.3% |

Figure 4 shows each tool’s annotation coverage for each genome using default settings (for *TRF* we used “`2 7 7 80 10 30` *<*max repeat period *>* `-l` 12”, as suggested in the TRF user guide). We also report results achieved with settings optimized for each organism; these settings were identified by performing a parameter grid search (see below). Finally, we show coverage for *ULTRA* using the “`--tune`” subprogram, which causes *ULTRA* to perform an automated grid search (described in Section Tuning).

**Figure 4.**
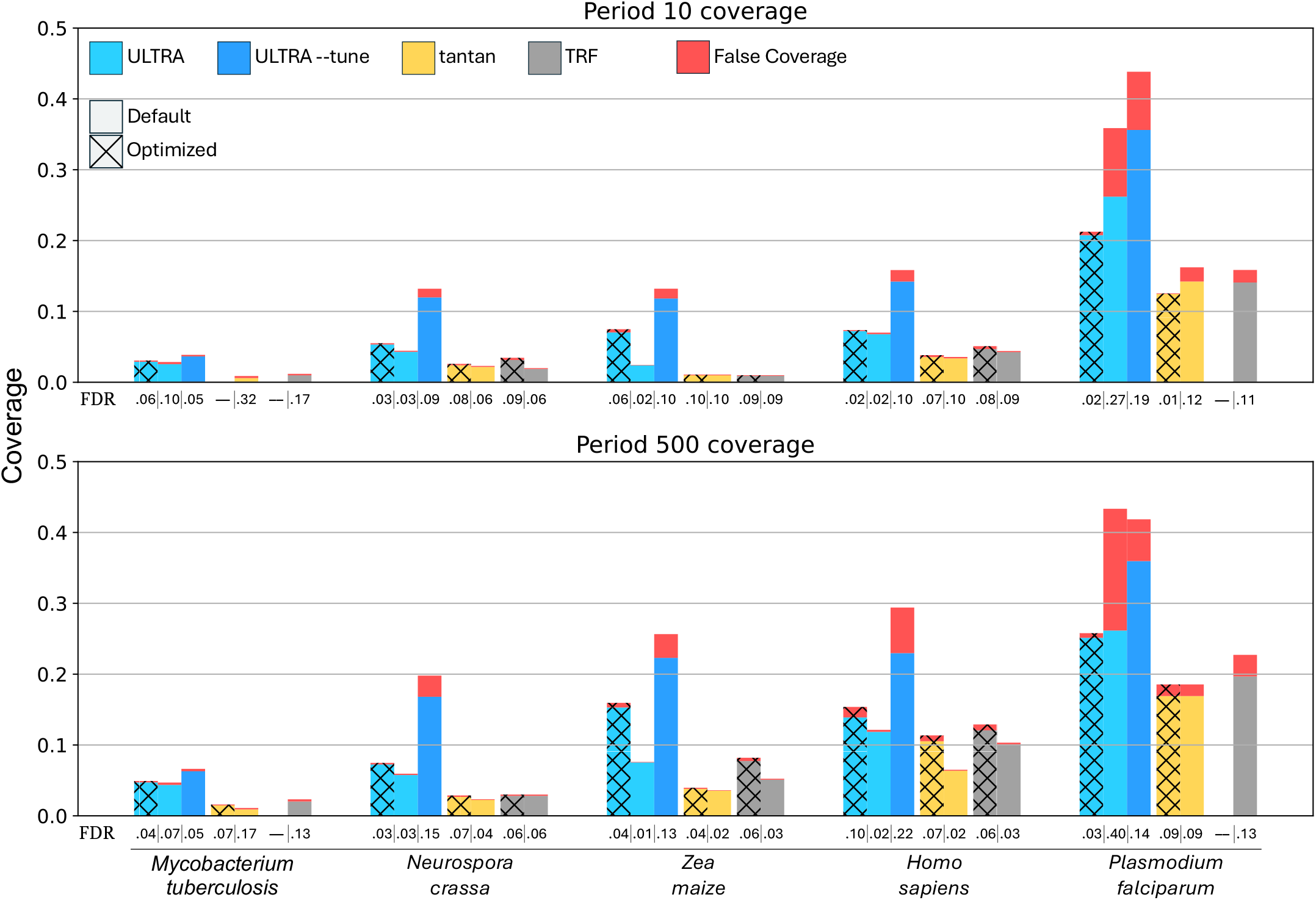
Coverage and estimated false coverage for *ULTRA, tantan*, and *TRF*. The top chart show coverage when using a maximum repeat period of 10 and the bottom chart show coverage when using a maximum repeat period of 500. Plain bars indicate default parameters and textured bars indicate grid-search optimized parameters. We also include results using *ULTRA* `--tune` (with default settings). The estimated false discovery rate (FDR) is displayed below each bar. Note that in some case, there is no parameter choice that achieves less than 10% FDR; in these cases, no bar is presented, and the FDR value is listed as —.

By default, *tantan* will annotate repetitive sequence even if there are fewer than two full repeat units, causing *tantan* to sometimes use a large repeat period to annotate noncontiguous repetitive content (i.e content that is not *tandemly* repetitive). To make *tantan*’s results more consistent with *ULTRA* and *TRF* we used “*tantan* `-f4`” for all coverage experiments, which (among other things) causes *tantan* to filter out repeat annotations that are less than two repetitive units in length. We also measured *tantan*’s performance without `-f4` and compare it against *ULTRA* with comparable settings (`--min unit` 0) and found that “*ULTRA* `--min unit` 0” has greater sensitivity than *tantan* without the `-f4` flag, with similar relative results.

For the optimized coverage experiments (Figure 4), we created a grid of parameters for each tool (see Table 2), and then for each genome, we selected the parameters that produced the highest coverage with an estimated false discovery rate of 10% or less.

**Table 2.**
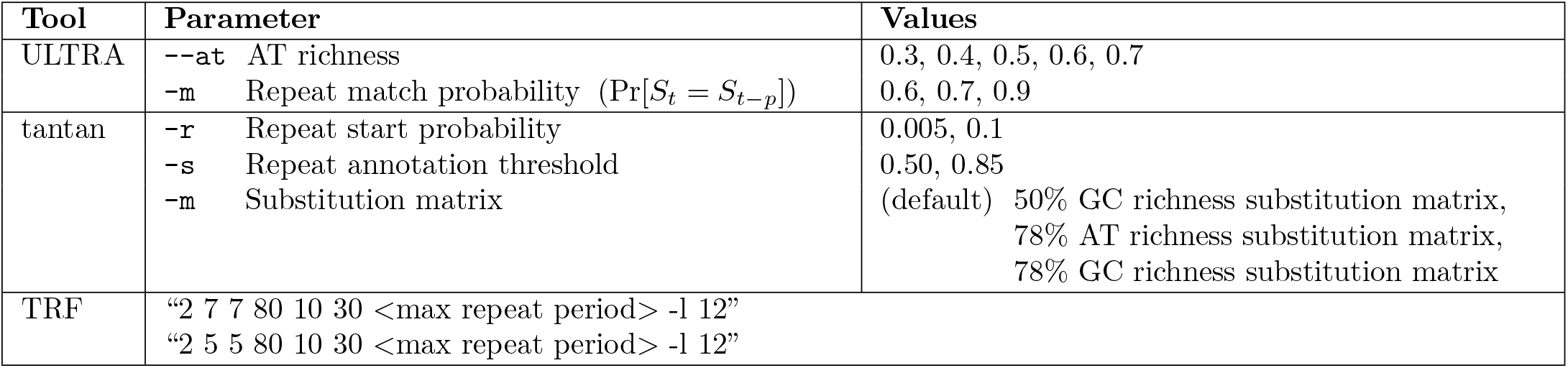
Parameter Grids for Optimized Coverage Experiments. *ULTRA*’s grid is composed of 15 total parameter combinations, *tantan*’s grid has 12 parameter combinations, and *TRF* grid has 2 different configurations, one from the TRF user guide, and one from [49].

| Tool | Parameter | Values |
| --- | --- | --- |
| ULTRA | --at AT richness | 0.3, 0.4, 0.5, 0.6, 0.7 |
| | -m Repeat match probability ($\Pr[S_t = S_{t-p}]$) | 0.6, 0.7, 0.9 |
| tantan | -r Repeat start probability | 0.005, 0.1 |
|  | -s Repeat annotation threshold | 0.50, 0.85 |
|  | -m Substitution matrix | (default) 50% GC richness substitution matrix,<br>78% AT richness substitution matrix,<br>78% GC richness substitution matrix |
| TRF | "2 7 7 80 10 30 <max repeat period> -l 12" |  |
|  | "2 5 5 80 10 30 <max repeat period> -l 12" |  |

For both default settings and optimized settings *ULTRA* produces greater coverage than *TRF* and *tantan*. For most genomes *ULTRA* also had the smallest FDR with the notable exception being *Plasmodium falciparum*, caused by the genome’s highly biased 80.7% AT-rich composition, which is far from *ULTRA*’s default composition expectation. For many genomes using *ULTRA* with `--tune` produced a larger FDR compared to *ULTRA* (default), especially in the period 500 coverage experiments. The increased FDR while tuning is primarily caused by `--tune` disabling indel states during tuning; for best results we suggest users use `--tune indel`.

### Repeat Score Distribution

When using repeat masking strategies, one must choose a score threshold that will determine which regions will and will not be masked. Understanding the distribution of annotation scores in random sequence allows researchers to make statistically informed decisions about which score threshold they should use, and may enable more sophisticated repeat-error mitigation than mere masking (such as competing annotations based on score density or associated statistical values [40]). Additionally, if a tool is incapable of producing well behaving score distributions when annotating random sequence, then it is likely that the tool will provide inconsistent scores to biological tandem repeats, resulting in lower quality repeat masking.

Figure 5 shows the score distributions of *ULTRA, tantan*, and *TRF* when run on 10 Gb of randomly generated 60% AT-rich sequence. We ran all three tools using default settings and a maximum detectable repeat period of 100. Its important to note that while *ULTRA* and *TRF* produce scores for each tandem repeat annotation, *tantan* instead produces probabilities (Forward-Backward derived posterior marginals) for individual letters.

**Figure 5.**
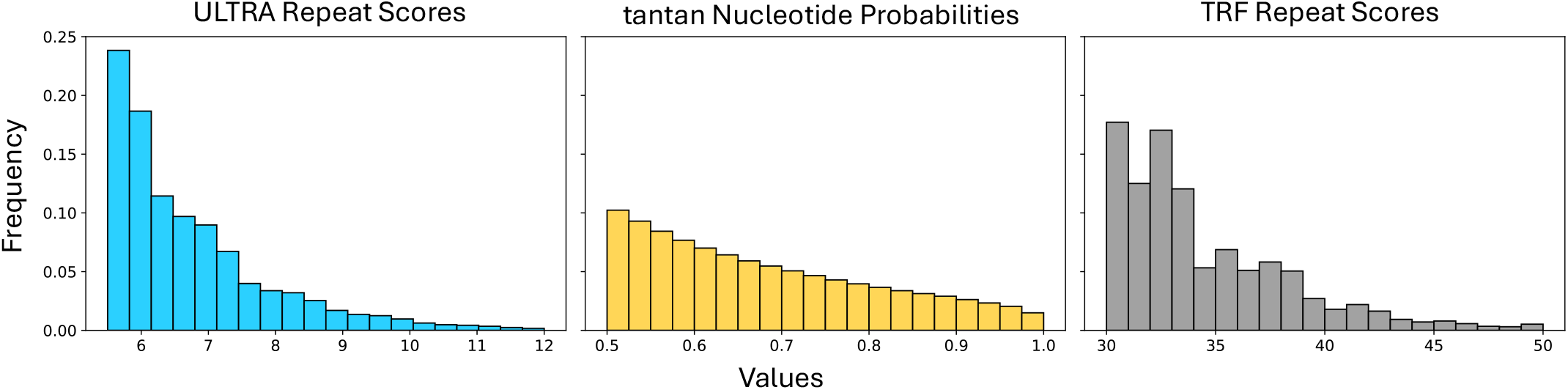
Annotation score distributions. Using each tool to label 10Gb of 60% AT-rich random sequence, the left and right plots show per-repeat score distributions for *ULTRA* and *TRF*, respectively. The middle plot shows the distribution of emphtantan per-letter probabilities of being part of a repetitive region. Horizontal axis corresponds to score/probability values and the vertical axis corresponds to value frequency. The exponential decay of ULTRA enables reliable P-value estiamtes.

*ULTRA*’s score distribution decays smoothly and exponentially, while *TRF* ‘s scores decay chaotically. *tantan*’s probability distribution cannot be compared directly to the scores produced by *ULTRA* or *TRF* because they represent letter-specific probabilities of being part of some tandem repeat instead of tandem repeat annotation scores, but we note that *tantan*’s posterior marginal distribution decays very smoothly.

### Repeat Splitting Accuracy

Repeat splitting (described in Section Repeat Splitting) allows *ULTRA* to find and annotate changes in repetitive pattern. Figure 6 shows *ULTRA*’s repeat splitting accuracy for repetitive periods 1 through 10 under substitution rates between 0 and 0.5. To measure repeat splitting accuracy, we generated artificial repetitive sequences that contained two subrepeats. Sequences were made by first selecting a repeat period between 1 and 10, and then creating two repeat units (such as “AACC” and “ACGT”) that shared no more than 50% similarity. The similarity restriction considers all rotations of the repetitive units; for example, “ACCGG” and “GGAGG” would *not* be a valid pair of subrepepeat patterns because “ACCGG” rotated two letters to the right is “GGACC”, which shares greater than 50% similarity with “GGAGG”.

**Figure 6.**
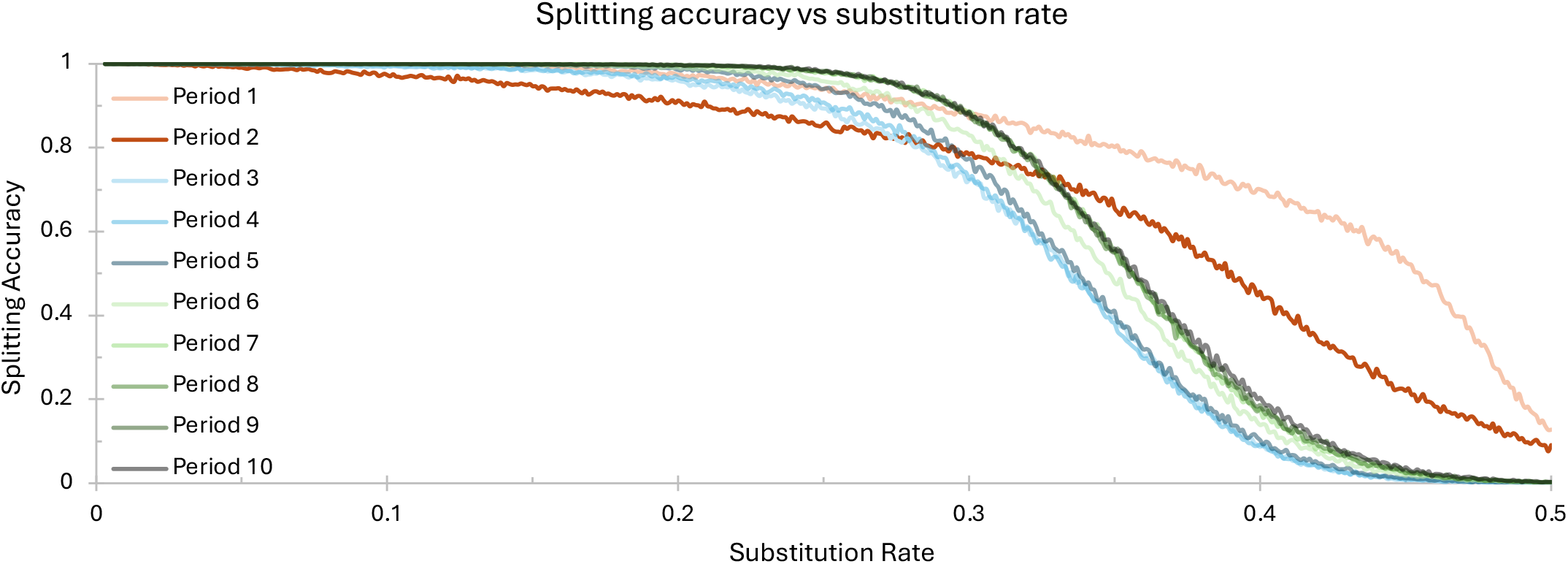
Repeat splitting accuracy vs sequence substitution rate.

We then generated two perfect tandem repeats of length 500, one for both repetitive units. We proceeded to concatenated the two perfect tandem repeats, creating one sequence of length 1000. We then mutated the sequence using a substitution rate ranging between 0 and 0.5, where each mutated letter is equally likely to mutate to any other letter.

For each combination of repeat period and substitution rate, we generated 5000 sequences and then ran *ULTRA* on each sequence. We quantify *ULTRA*’s repeat splitting accuracy as the fraction of sequences in which *ULTRA* found a single change in repetitive pattern that was within 10 nucleotides of the true change in repeat pattern. Figure 6 demonstrates that *ULTRA*’s repeat splitting is generally accurate for repetitive sequence with a substitution rate of 30% or less. This suggesting that it may be appropriate to report repeat units for subrepeats estimated to contain sequence identity *>* 70%, and to otherwise report a core “ambiguous” repeat unit for the entire block.

### Computational Resources

Table 3 shows memory usage and runtime while processing the T2T genome (3.2 gb) with *ULTRA, TRF*, and *tantan*. All analysis was performed on 2.4GHz XE2242 CPUs with 94 cores and 512 GB of RAM. *TRF* with a maximum detectable repeat period of 10 failed to process all T2T chromosomes without crashing despite efforts to adjust the “-l” option and increase *TRF* ‘s allocated memory. *ULTRA* is the only tool capable of multithreading, and all of *ULTRA*’s analysis was performed using 16 threads. *ULTRA*’s memory footprint can linearly be decreased by reducing the number of threads used.

**Table 3.** Computational time/memory used while annotating the T2T genome using. Default settings were used for *ULTRA*_10_, *TRF*, and *tantan. ULTRA*_5_ uses 5 insertion/deletion states, and *ULTRA*_0_ uses no insertion/deletion states. *TRF* (10) crashed before annotating all chromosomes.

| Tool (period) | User Time (Hours) | Wall Time (Hours) | Peak Memory Usage (GB) |
| --- | --- | --- | --- |
| *ULTRA*<sub>10</sub> (10) | 1.20 | 0.08 | 1.1 |
| *ULTRA*<sub>5</sub> (10) | 0.89 | 0.06 | 1.0 |
| *ULTRA*<sub>0</sub> (10) | 0.16 | 0.02 | 0.7 |
| *tantan* (10) | 0.02 | 0.02 | 1.3 |
| *TRF* (10) | 8.66 | 8.66 | 6.1 |
| *ULTRA*<sub>10</sub> (500) | 114.83 | 7.17 | 3.7 |
| *ULTRA*<sub>5</sub> (500) | 60.80 | 3.82 | 2.1 |
| *ULTRA*<sub>0</sub> (500) | 6.50 | 0.40 | 2.0 |
| *tantan* (500) | 0.56 | 0.56 | 1.3 |
| *TRF* (500) | 10.20 | 10.20 | 19.6 |

## Discussion

Here, we have introduced our open-source tool for identifying and labeling repetitive DNA: *ULTRA* (https://github.com/TravisWheelerLab/ULTRA), in hopes that it both aids the study of tandem repeats and also helps reduce the bioinformatic problems caused by tandem repeats. A prototype implementation of *ULTRA* was used during the development of T2T-CHM13 [31] and the 1.0 version of *ULTRA* will soon be integrated into *RepeatMasker* [41] (replacing *TRF*). We plan to continue supporting and improving *ULTRA* and have already strategized algorithmic tweaks and model changes that will further increase *ULTRA*’s speed, sensitivity, and specificity. We also highlight that *ULTRA* 1.0 supports only DNA and RNA sequences; support for protein sequences will be added in a future release.

The patterns of tandem repeats are complex and far from fully understood. Higher order repeats are particularly complex, and there is great need for methods that characterize their hierarchical structure and repetitive patterns. As we continue supporting and developing *ULTRA* we hope to expand upon the descriptive annotations currently provided by *ULTRA* so that repetitive sequence can be better characterized and better understood.

## 1 Acknowledgments

We thank Robert Hubley, Arian Smit, and Tim Anderson for helpful discussions during development of software and benchmarks. We also gratefully acknowledge the computational resources and expert administration provided by the University of Montana’s Griz Shared Computing Cluster (GSCC), CyVerse’s External Collaborative Partnership program (provided by Tyson Swetnam), and the high performance computing (HPC) resources supported by the University of Arizona TRIF, UITS, and Research, Innovation, and Impact (RII) and maintained by the UArizona Research Technologies department.

## 2 Funding

The authors were supported by NIH NIGMS grants P20GM103546 and R01GM132600. DRO was additionally supported by the National Institute of Allergy and Infectious Diseases (NIAID), National Institutes of Health (NIH), Department of Health and Human Services under BCBB.

## Notes

### Competing Interest Statement

The authors have declared no competing interest.

